# A Key Regulator of Dendritic Morphology in Supragranular Neocortex Impacts Mismatch Negativity

**DOI:** 10.64898/2026.01.28.702244

**Authors:** Anna M. Rader Groves, Connor G. Gallimore, Valentina J. Sutton, Jordan M. Ross, Robert A. Sweet, Melanie J. Grubisha, Jordan P. Hamm

## Abstract

**Background:** Cytoskeletal structure and neuronal function are heavily intertwined, and altered pyramidal neuron morphology (e.g., reduced dendritic arbors) has been consistently identified in the neocortex of individuals with psychiatric disorders. A missense mutation in *Kalrn* (*Kalrn*-PT) enhances activation of RhoA, a cytoskeletal regulator, and leads to adolescent-onset dendritic regression in supragranular auditory cortex. To investigate the relationship between this spatiotemporally altered cytoskeletal structure and psychiatric-relevant dysfunction, we examine the functional impacts of this genetic mutation, focusing on mismatch negativity – a common biomarker of altered sensory integration that matures across adolescence.

**Methods:** In *Kalrn*-PT and littermate wild-type mice, pyramidal dendritic morphology was assessed across primary visual cortex (V1) layers using Golgi staining (n=172 neurons/26 mice). Local field potentials (LFPs) and neuronal spiking were recorded during a visual oddball sequence (n=25 mice). Deviance detection – the rodent neural analogue of mismatch negativity – was analyzed across V1 layers.

**Results:** *Kalrn*-PT mice exhibit reductions in V1 dendritic length specific to layer 2/3, mirroring observations in post-mortem samples from people with psychiatric disorders. Deviance detection was reduced in LFPs from *Kalrn*-PT mice and absent in layer 2/3 neurons, despite typical feature selectivity and firing rates. These deficits were accompanied by reduced functional connectivity between visual and frontal areas.

**Conclusions:** These results highlight alterations in higher-level sensory cortical integration in *Kalrn*-PT mice and demonstrate that adolescent-onset, disease-relevant structural and functional phenotypes can be linked by a common upstream effector. Further, these alterations appear restricted to supragranular layers, demonstrating an outsized role for *Kalrn*, particularly longer isoforms, in superficial neocortex.

## Introduction

The structure of neurons is tightly linked to their function and physiology, and thus, pathological neuronal morphology is often connected to pathological neuronal function (1,2). In neuropsychiatric disorders such as schizophrenia and bipolar disorder, atypical neuronal morphology, including reductions in dendritic arbors and dendritic spines, is commonly observed postmortem (3). Such alterations have been reported in several regions, including dorsolateral prefrontal cortex, primary auditory cortex (A1), and primary visual cortex (V1) (4–6). Notably, morphological abnormalities have been largely observed in supragranular layers (layer 2/3) (7–9), which aligns with clinical genetic risk and gene expression data implicating supragranular neurons in schizophrenia etiology and suggests a pathological role for these supragranular abnormalities (10,11).

A missense mutation (P2255T) in the *Kalrn* gene, *Kalrn*-PT, has been shown to reduce dendritic length and complexity in layer 3 (but not layer 5) of mouse A1, recapitulating a commonly reported finding in postmortem cortical tissue from individuals with schizophrenia (6,12,13). *KALRN*-PT was first identified as a rare risk-modifying variant in a Japanese population with schizophrenia (14), although a subsequent large-scale examination of schizophrenia-associated mutations did not identify *KALRN* ((15) but see also (16)). The mutation, which is only present in longer isoforms (Kal9, Kal12), directly influences dendritic structure through a RhoA gain of function, a molecular pathway with established clinical relevance (17–22). Importantly, altered pyramidal arbors in mice with the *Kalrn*-PT variant are present in adulthood but not before adolescence, mirroring the adolescent onset of neuropsychiatric disorders like schizophrenia (12).

As many have emphasized (23–25), no mouse model can replicate complex, polygenic, polyetiological psychiatric diseases like schizophrenia with broad validity. Instead, models should focus on a restricted set of disease-relevant domains. To this end, *Kalrn*-PT mice offer a narrow model of molecular (RhoA) and cellular (dendrite loss) alterations with schizophrenia-relevant spatial (supragranular) and temporal (adolescent-onset) features arising from a single genetic (base pair) change (8,12,17,26). By recapitulating these facets, *Kalrn*-PT mice provide a controlled system for linking such molecular and structural alterations to cell and circuit functions. Exploring functional alterations that co-occur with these morphological deficits in clearly defined animal models helps delineate clusters of common psychiatric features sensitive to – and linked by – a common upstream effector.

Reduced mismatch negativity (MMN), an electrophysiological measure indexing sensory context processing (i.e., heightened responses to unexpected stimuli), is one of the most consistently reported biomarkers of schizophrenia (27). Altered MMN offers a candidate biomarker that may be impacted by the *Kalrn*-PT mutation for several reasons. 1) Mismatch responses in populations with schizophrenia have been correlated with gray matter volume reductions thought to reflect the loss of neuropil (including dendrites), but direct measures of dendritic structure are not accessible in humans *in vivo* (28–30). 2) Past research indicates that deviance detection (a rodent analogue of MMN) is first computed in supragranular pyramidal neurons of primary sensory cortex, the aberrant cell type observed in *Kalrn*-PT mice (31,32). In supragranular neurons, bottom-up sensory afferents are integrated with top-down feedback from higher brain regions (33,34). Our past work (31,35) and others’ (36,37) have demonstrated that such feedback input to sensory cortices is required for deviance detection. It is reasonable then to expect deviance detection to rely on intact dendritic structure in layer 2/3 neurons themselves, as these arbors provide the substrate for distributed input integration (38). 3) In humans, MMN amplitude and topography have been shown to mature during adolescence (39–41). Critically, work from our group has recently shown that deviance detection in primary sensory cortex emerges across adolescence in mice as well (42). Thus, deviance detection relies on the specific computational units (i.e., supragranular neurons) impacted in *Kalrn*-PT mice *and* follows a developmental trajectory common to the model’s morphological phenotype and schizophrenia symptom onset.

Using the *Kalrn*-PT model, we sought to test the hypothesis that spatially and temporally selective cytoskeletal structural alterations co-occur with altered sensory context integration. To do so, we evaluated integrative sensory measures, including deviance detection and orientation selectivity, in *Kalrn*-PT mice across cortical layers, along with pyramidal neuronal function. Together, the results address a fundamental gap in understanding how dendritic structure relates to visual cortical function and indicate a role for *Kalrn* in sensory physiology. Moreover, the circumscribed nature of the alterations in supragranular layers present a plausible link from mechanism to phenotype.

## Methods and Materials

All experimental procedures were carried out under the guidance and supervision of the Georgia State University (GSU) Division of Animal Resources and in accordance with the approved Institutional Animal Care and Use Committee (IACUC) protocol at GSU. See Supplemental Methods for detailed explanations.

### Animals

Adult (12 weeks or older) female and male mice were used for these experiments. Mice were generated from breeding pairs heterozygous for the P->T mutation (P2255T) in the *Kalrn* gene, permitting use of *Kalrn*-PT homozygous and wild-type (WT) littermates. These mice were previously generated on a C57BL/6 background using CRISPR/Cas9, validated (e.g., assessed for off-target effects), and shown to have unaltered expression of major Kalirin isoforms (Kal7, Kal9, Kal12) (12).

### Golgi Histology and Imaging

V1 sections were taken from *Kalrn*-PT and littermate/cagemate WT mice and stained according to the FD Rapid Golgi Stain Kit protocol (FDNeurotechnologies). Golgi-stained tissue from V1 of WT (n=13 mice, 85 neurons) and *Kalrn*-PT mice (n=13 mice, 87 neurons) was imaged across layers 2/3-5. Only basilar arbors were examined for simplicity given the similar reductions between basilar and apical arbors in A1 of *Kalrn*-PT mice and because basilar measures are more commonly reported (12,43). The selection criteria for neurons included: non-overlapping soma, complete staining (i.e., no truncated branches), and at least three primary basilar dendrites.

### Electrophysiology and Visual Stimuli

Electrophysiological data consisted of two main cohorts generated on independent experimental set-ups (Fig. 2B): Cohort 1, bipolar electrodes implanted in V1 and anterior cingulate area (ACa), 12-18 weeks, n=8 WT (3 female) and n=8 *Kalrn*-PT (4 female); Cohort 2, multielectrode spanning 750 μm across V1 layers (dorsal/ventral axis), 18-20 weeks, n=4 WT (3 female) and n=5 *Kalrn*-PT (3 female). Recordings were conducted during two types of visual sequences. First, a “many-standards” control sequence was shown, made up of 8 distinct orientations presented randomly with approximately equal likelihood (≈12.5% probability). Next, an oddball sequence was displayed, composed of two stimuli separated by 90-deg (e.g., 0-deg and 90-deg), one shown in a redundant context (≈87.5% probability) and one in a deviant context (≈12.5%) presentation. Contexts were “flip-flopped” within the same run, halfway through, such that the previously redundant orientation served as the deviant and vice versa. Stimuli were presented at 100% contrast, 0.8 cycle per degree, 2 cycles per second.

This procedure ensured that we captured responses to a given orientation in all three sensory contexts: equiprobable (control), high likelihood/expected (redundant), and low likelihood/unexpected (deviant).

### Analysis

#### Histology

To assess genotype differences in dendritic morphology, the area under the curve across the 30-70um Sholl radii (proximal AUC) was taken for dendritic length and number of Sholl intersections, as proximal dendrites showed the greatest genotype difference qualitatively. Linear mixed effects modelling and estimated marginal means were used to test for within-layer effects while controlling for intra-mouse correlations across cells. One-sided testing was used given our *a priori* expectations based on previous observations in *Kalrn*-PT A1.

#### Electrophysiology

To compare stimulus-induced power, we combined single trial power estimates across both cohorts (single and multielectrode contexts; n=12 WT, n=13 *Kalrn*-PT) and focused on theta to low-gamma power (4-60Hz), averaged from 40 to 350ms post-stimulus onset and across orientations. We carried out a linear mixed effects analysis with context as a within-subjects variable and genotype and sex as between-subjects variables. Mouse and cohort (single vs multielectrode recordings) were treated as a random effects variable.

ACa-to-V1 coherence (Cohort 1; n=7 WT, n=8 *Kalrn*-PT) was analyzed using a mean-vector technique on the ACa–V1 phase differences – similar to an inter-trial phase coherence metric (44) as previously described (35). One WT animal was removed due to missed ACa electrode placement. Statistical analysis focused on the peak in synchrony between 6 and 10HZ, and a mixed ANOVA assessed genotype and stimulus context.

For analysis of neuronal spiking, we analyzed single unit (n=83) and multiunit cluster activity (n=104) in multielectrode recordings (Cohort 2; n=187 from 3 WT and 5 *Kalrn*-PT animals). One WT animal was excluded because units were lost between recordings. Stable units (see supplemental methods for criteria, Supplemental Fig. 2) were split into broad spiking (Fig. 3) and narrow spiking units/clusters (Supplemental Fig. 4). For each unit, spikes were averaged across trials for each stimulus context type (control, deviant). For statistical analyses of deviance detection, we focused on the early stimulus evoked peak (occurring in the first 150ms post stimulus onset) and the average sustained activity (150ms-350ms) separately. We carried out linear mixed effects analysis on individual unit response metrics (deviance detection early peak, sustained activity, SSA, OSI, global firing rate) with mouse as a random effects variable. We split analyses into three putative layers: layer 2/3 (200 to 350µm), layer 4(351-450µm), and layer 5/6 (451-750µm).

## Results

### Layer 2/3 pyramidal neurons exhibit reduced dendritic length in *Kalrn*-PT V1

Decreased dendritic arborization in pyramidal neurons, particularly in supragranular layers, has been found across several cortical regions (e.g., V1, prefrontal cortex) in schizophrenia (6,13). A recent study by our group showed that *Kalrn*-PT mice recapitulate these deficits in layer 3 (but not layer 5) of A1 (12). We anticipated that these deficits would extend to additional sensory cortical regions and thus be present in V1. To test this prediction and further assess the laminar stratification of these effects, samples from V1 of *Kalrn*-PT and WT littermates were Golgi-stained, and basilar dendritic morphology of pyramidal neurons was compared between genotypes for layer 2/3, layer 4, and layer 5 (Fig. 1). To minimize statistical tests, we focused analyses on the area under the curve (AUC) for proximal branches (30-70um radii), as this captured the peaks.

**Figure 1.**
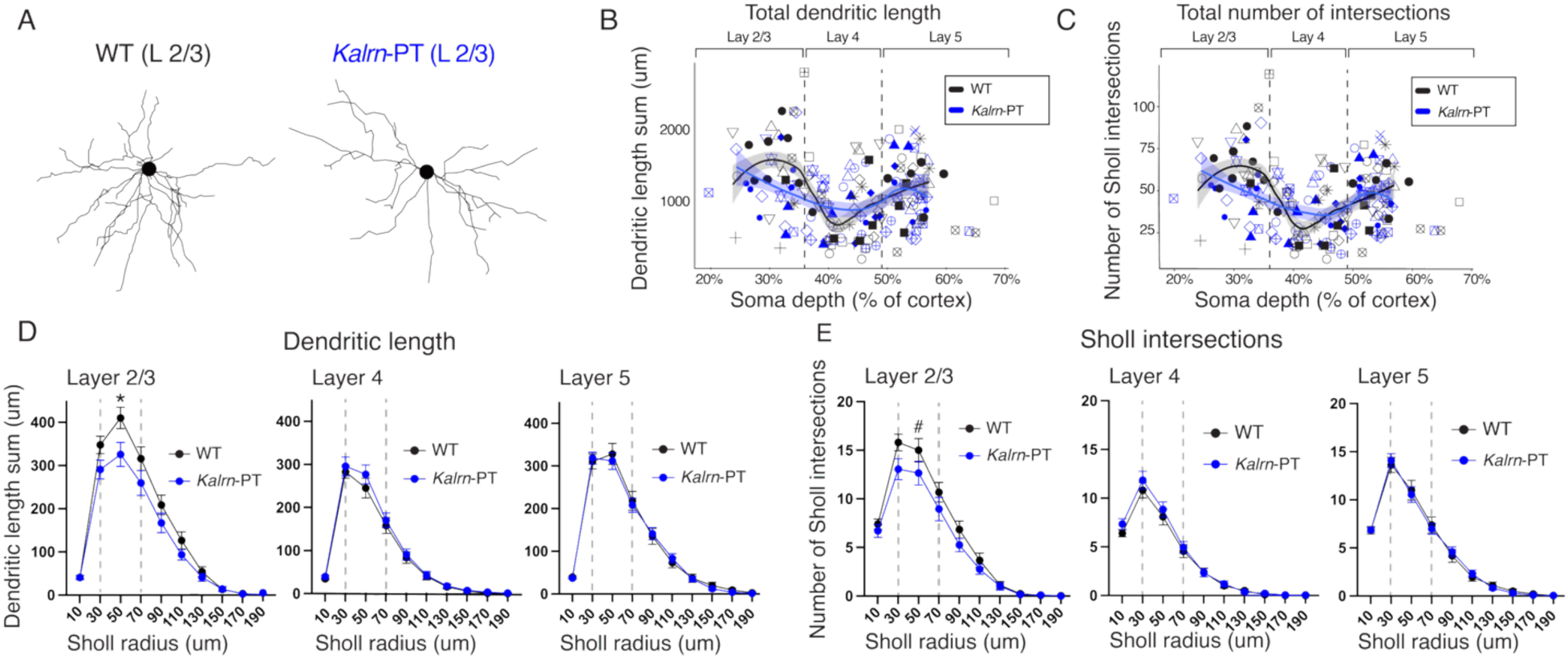
*Kalrn*-PT mutation decreases V1 proximal dendritic arbors in layer 2/3. **(A)** Representative reconstructions of basilar dendritic arbors from pyramidal neurons in V1 layer 2/3 from WT (left) and *Kalrn*-PT (right) animals. **(B)** Total length of the basilar dendritic arbor and **(C)** total number of basilar Sholl intersections as a function of soma depth in V1 pyramidal neurons from WT animals (n=85 cells, 13 animals) and *Kalrn*-PT (n=87 cells, 13 animals). Lines showing LOESS smoothed fit (span=0.6) are overlaid with shaded regions indicating SEM. Matching shapes indicate cells from the same animal. **(D)** Sum of dendritic length and **(E)** number of Sholl intersections by Sholl radius in layer 2/3 (WT: n=25 cells/12 animals; *Kalrn*-PT: 20 cells/8 animals), layer 4 (WT: n=36 cells/12 animals; *Kalrn*-PT: n=32 cells/12 animals), and layer 5 (WT: n=24 cells/11 animals; *Kalrn*-PT: 35 cells/13 animals). In D-E, error bars indicate SEM. * indicates p < 0.05. # indicates trend-level significance (p = 0.058). SEM = standard error of the mean; WT = wild type.

Proximal dendritic length differed significantly between genotypes only in superficial layers (Fig. 1D; Layer 2/3: t(87.2)=1.985, p=0.0251; Layer 4: t(65.1)= -0.790, p=0.7838; Layer 5: t(82.6)=0.227; p=0.4104). For proximal dendritic complexity measures (i.e., number of Sholl intersections), superficial layers exhibited a trend towards genotype differences, but this difference did not reach statistical significance (Fig. 1E; Layer 2/3: t(86.2)=1.588, p=0.058). Layers 4 and 5 showed no differences in complexity by genotype (Fig. 1E; Layer 4: t(61.2)= - 0.620, p=0.7313; Layer 5: t(76.2)=0.012, p=0.4954). These analyses revealed a decrease in the extent of proximal dendritic arbors in supragranular, but not granular or infragranular, pyramidal neurons in V1 (Fig. 1D-E), consistent with previous data in A1 of *Kalrn*-PT mice and in patterns of spine density alterations in postmortem SZ tissue (8,12).

In addition to dendritic arbors, dendritic spine density is often reported to be decreased in schizophrenia (4,43). Spine density per unit tissue was previously shown to be reduced in layer 3 *Kalrn*-PT A1, but spine density per unit dendrite length was reported unaltered, suggesting that this decrease in spine tissue density is driven by the decrease in dendritic length (12). In a smaller cohort, we observed a trend towards decreased spine density (i.e., PSD-95 puncta density) in superficial V1 of *Kalrn*-PT mice (Supplemental Fig. 1), consistent with our observations of reduced dendritic arbors within this region.

### Visual deviance detection, the intracortical rodent analogue of mismatch negativity, is abolished in *Kalrn*-PT mice

Given reductions in supragranular dendrites in V1, we next sought to assess a functional measure of cortical integration with disease relevance, focusing specifically on visual deviance detection during an oddball sequence. Deficits in visual mismatch negativity, though frequently overlooked in favor of the auditory domain, are consistently reported in schizophrenia and other disorders of sensory-perceptual dysfunction (45,46). Other visual perceptual aberrations are commonly reported in psychiatric disorders as well, including in early phases of schizophrenia (47–49).

We recorded extracellular electrophysiology from V1 in awake WT and *Kalrn*-PT mice using either a bipolar electrode (n=16) or a multielectrode array (n=9; Fig. 2B). During electrophysiological recordings, mice were shown a standard visual oddball sequence (repetitive stimuli randomly interrupted by a “deviant” stimulus) and a many-standards control sequence (stimuli are equiprobable) (Fig. 2A). Stimulus-induced power was measured for two stimulus contexts: a) contextually neutral (i.e., during a many-standards control) vs b) contextually deviant (“oddballs”). The difference in responses to these two stimulus contexts equates to deviance detection. Using these two separate sequences (instead of comparing between the deviant and the repeated “redundant” stimulus within the oddball run) is necessary to differentiate cortically dependent “deviance detection” from stimulus specific adaptation (32), which has been previously identified to emerge subcortically (50). A linear mixed effects analysis revealed a genotype-by-stimulus context interaction (t(42)= -2.4115, p=0.020), indicating a significant reduction in deviance detection in *Kalrn*-PT mice (Fig. 2C-E).

**Figure 2.**
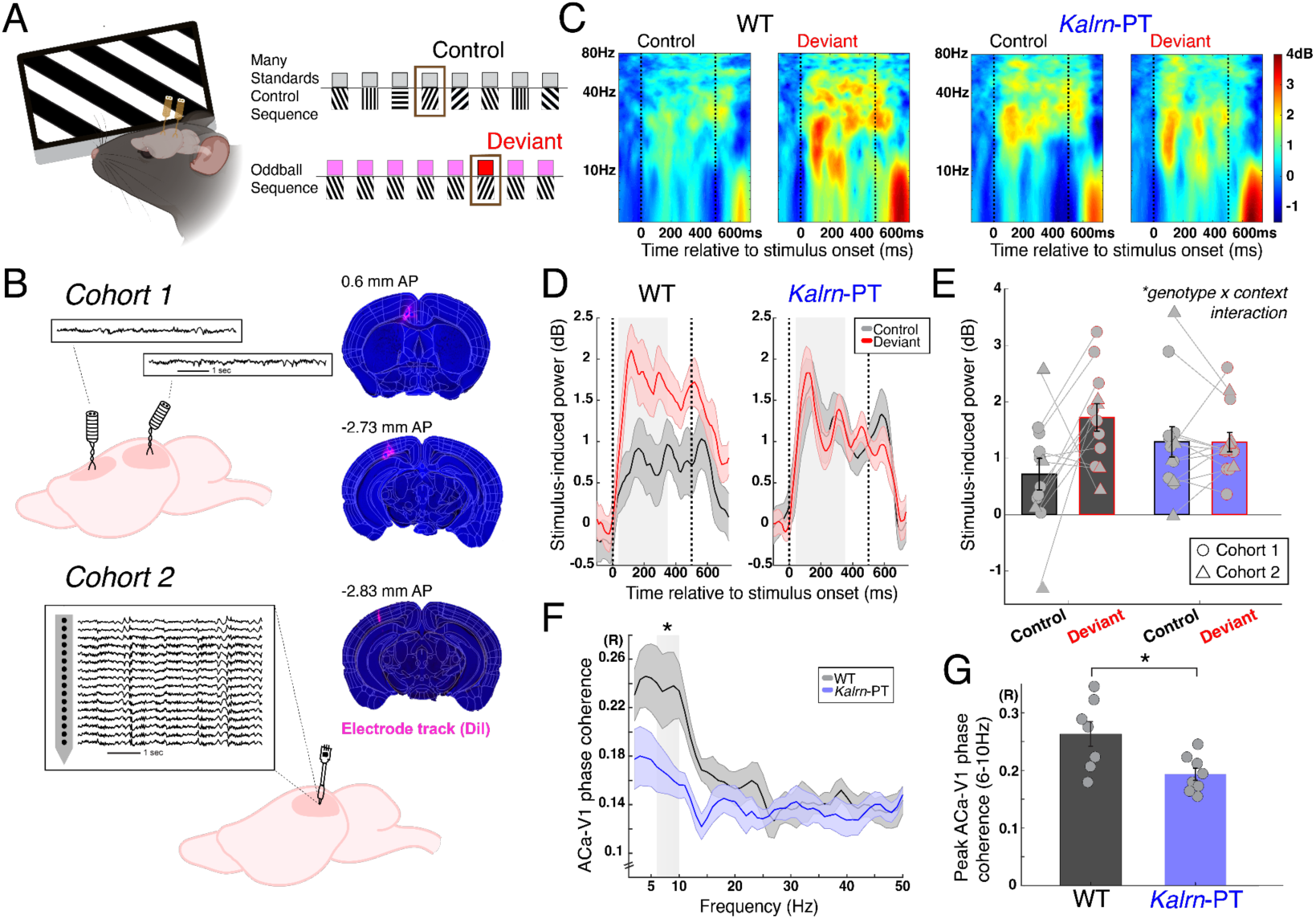
*Kalrn*-PT mice exhibit reduced deviance detection and fronto-visual synchrony. **(A)** Awake mice viewed full field moving gratings. Visual stimuli include the many-standards control sequence (top) in which eight stimuli are equiprobable and the oddball sequence (bottom) in which one orientation is repeatedly shown and interrupted by a different (deviant) orientation randomly. Brown box marks the same stimulus in the control and deviant context. **(B)** Local field potentials were recorded from two cohorts. *Cohort 1 (top)*: Bipolar electrode recordings were taken from V1 and ACa. *Cohort 2 (bottom)*: 16-channel multielectrode recordings (750 µm span, 50µm interelectrode spacing, from layer 1 to 5) were taken from V1. Electrode placement was confirmed with histology following recordings in both cohorts. **(C)** Stimulus-induced power in V1 during control and deviant stimulus contexts for WT (left) and *Kalrn*-PT (right) animals. **(D)** Stimulus-induced power (averaged 4-60Hz) by context across the stimulus presentation interval for WT (left) and *Kalrn*-PT (right) mice (± SEM). **(E)** Average stimulus-induced power (± SEM) by context from (D; 40-350ms) for WT (left) and *Kalrn*-PT (right) mice. Individual mice from Cohort 1 and 2 are overlaid as circles and triangles, respectively. Panels C-E combine data from Cohorts 1 and 2. **(F)** ACa-V1 phase coherence (± SEM) during visual stimulation, averaged across all stimulus conditions for WT (n=7) and *Kalrn*-PT (n=8) animals. **(G)** Peak ACa-V1 phase coherence across theta-band frequencies (6-10Hz) per animal. Panels F-G include data from Cohort 1 only. * indicates p < 0.05. SEM = standard error of the mean; WT = wild type. Dashed lines indicate stimulus onset/offset and shaded boxes indicate time or frequency window used for statistical analysis.

We have previously shown that deviance detection responses in V1 depend on feedback from ACa (a prefrontal region that projects to V1 in mice), particularly in the theta-alpha band (31,35). To assess the integrity of fronto-visual circuitry, we measured phase coherence between local field potential (LFP) signals in ACa and V1. The theta band (6-10Hz) peak in ACa-V1 synchrony observed in WT mice was significantly reduced in *Kalrn*-PT mice during visual stimulation (main effect of genotype, F(1,13) =7.37, p=0.018), suggesting impaired inter-regional communication (Fig. 2F-G).

### Neural responses to sensory mismatch are impaired in layer 2/3 in *Kalrn*-PT mice

To test layer-specific context integration in *Kalrn*-PT mice, we isolated spiking units (putative single neurons or local multineuron clusters) stable across multielectrode recordings in Cohort 2 (n=187 from 5 *Kalrn*-PT mice and 3 WT mice; Supplemental Fig. 2) and analyzed activity during visual stimuli. To reflect the cell type in which we observed altered morphology, we focused on units that exhibited broad spike waveforms characteristic of pyramidal neurons (n=147) for the main analysis (see Supplemental Fig. 4 for narrow spiking units).

In WT mice, we found robust deviance detection in putative layer 2/3 neurons that emerged early, <100ms after the onset of the stimulus (Fig. 3A,B). In contrast, layer 2/3 neurons in *Kalrn*-PT mice did not show strong deviance detection in this period (Fig. 3C,D). A linear mixed effects analysis, with mouse as a random effects variable, confirmed this genotype-by-context interaction on peak firing rate from 0 to 100ms (significant fixed effects coefficient; t(76)=-2.80, p=0.006; Fig. 3E) within layer 2/3. This effect was driven by units spread evenly across mice and was not driven by one recording (Fig. 3E). This finding aligns with the LFP data (Fig. 2C-E) and, as a measure of integrated neuronal output, further suggests that supragranular pyramidal cells fail to propagate deviance detection responses.

**Figure 3.**
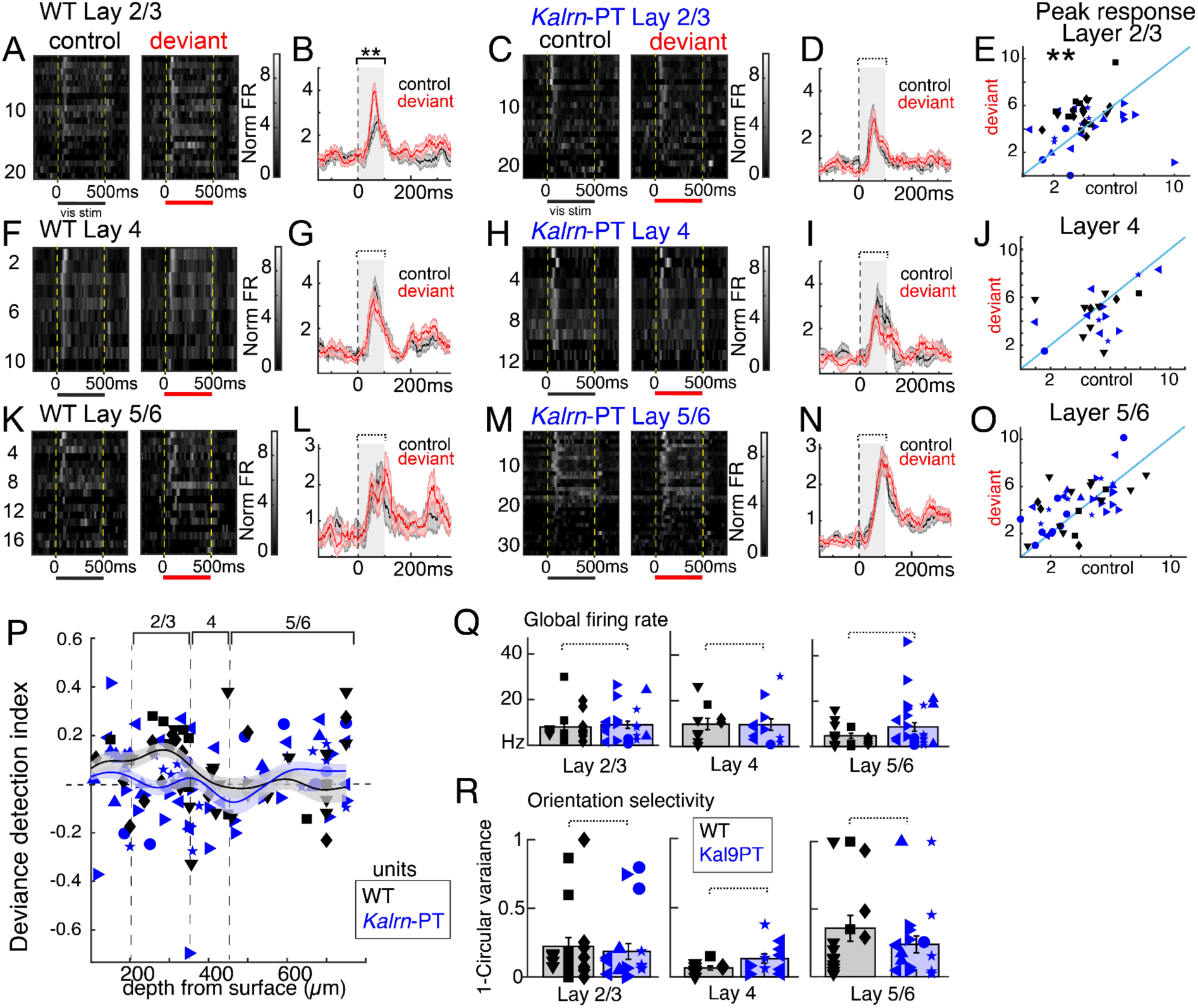
Layer 2/3 neurons fail to display deviance detection in *Kalrn*-PT mice. **(A)** Raster plots of trial-averaged firing responses (normalized firing rate; FR) across units (putative single and multineuron clusters) from layer 2/3 from WT mice. **(B)** Responses from (A) averaged over units with shaded regions indicating SEM. **(C-D)** same as A-B, but for *Kalrn*-PT mice. **(E)** Peak responses from 0 to 100ms after the onset of stimuli. Each dot is one unit. **(F-J)** Same as A-E for layer 4 neurons. **(K-O)** Same as A-E for layer 5/6 neurons. **(P)** Deviance detection index (deviant minus control, divided by max response over all contexts) plotted as a function of depth from pial surface. Smoothed gaussian fit line (200µm window) overlaid for each group. **(Q)** Overall firing rates across recordings and **(R)** orientation selectivity (1 minus circular variance) did not differ between genotypes for any layer. For E-R, matching shapes indicate isolated units from the same animal. ** p<0.01, genotype-by-context interaction. Dashed lines indicate p>0.05. SEM = standard error of the mean; WT = wild type.

We did not identify clear deviance detection, genotype differences, or genotype-by-context interactions for early or late latency responses in putative layer 4 (Fig. 3F-J) or layer 5/6 units (Fig. 3K-O). To further summarize how deviance detection effects were spread across cortical depth, we generated a “deviance detection index” by subtracting the deviant response from the control and then dividing this by the maximum response across all stimulus conditions for each unit. These values are plotted in Fig. 3P.

### Basic visual processing remains intact in *Kalrn*-PT mice

To verify that mismatch deficits are not simply a consequence of gross alterations to neuronal function, we examined global firing rate and orientation selectivity. We found unaltered firing rate (Fig. 3Q), and normal orientation selectivity across units in layer 2/3 (Fig. 3R; 1-circ variance; t(33)=1.10, p=0.28), layer 4 (t(18)=1.72, p=0.10), and layer 5/6 (t(91)=0.67,p=0.50) across these same broad spiking units (putative pyramidal neurons). Moreover, we observed normal visually evoked responses in *Kalrn*-PT mice (i.e., no genotype main effects in any layer, see e.g., Fig. 3A vs 3C).

Stimulus specific adaptation – reduced responses to repeated stimuli of the same orientation, relative to the control run – was also not altered in any layer in *Kalrn*-PT mice (Supplemental Fig. 3). Previous reports suggest that deviance detection, not stimulus-specific adaptation, is the component of mismatch negativity altered in schizophrenia, suggesting that *Kalrn*-PT mice model the specific mismatch deficits common in the disorder (51).

Further, narrow spiking units (putative parvalbumin interneurons) did not exhibit dramatic deviance detection in either group and were not grossly altered in firing rate or feature selectivity in any layer (Supplemental Fig. 4).

In summary, just as dendritic length was decreased in *Kalrn*-PT mice specifically in layer 2/3, we found that neural deviance detection – which is adolescent emergent and computed in layer 2/3 – was abolished in *Kalrn*-PT mice, and confirmed that this altered activity was specific to layer 2/3 neurons. This was against the backdrop of otherwise typical visual processing, suggesting that basic properties are spared in this model while more complex and emergent aspects of sensory processing are affected.

## Discussion

In complex, polygenic, and polyetiological diseases like schizophrenia, targeted study of limited (but well-delineated) genetic/molecular risk states in mouse models can isolate convergence points and upstream mechanisms for precision psychiatry. The data presented here illustrate how a missense mutation – a naturally occurring *Kalrn* variant – can lead to coinciding functional and structural neuropathology, specifically in visual cortex, an often-overlooked site in translational work despite its relevance in early schizophrenia (47,49,52,53)

While *Kalrn*-PT is not a highly penetrant risk gene for schizophrenia, several studies support a broader role for the *KALRN*-dependent regulation of RhoA or related pathways in schizophrenia pathology (17). For example, elevated Kal9 levels in sensory cortex (which lead to reduced dendritic arbors *in vitro*) were observed in 77% of individuals with schizophrenia in one cohort (54). In a separate cohort with psychiatric disorders, multiple molecular signatures of RhoA overactivation (e.g., increased PDZ-RhoGEF, increased phospho-cofilin) were observed postmortem in dorsolateral prefrontal cortex (18). Decreased actin polymerization (i.e., F-actin ratio) and altered expression profiles of other related genes (e.g., GTPases, GEFs, and myelin-associated inhibitors (MAIs)) have also been reported (55–57).

Related genetic mutations further implicate *KALRN*-related pathways in the etiology(ies) of psychiatric disorders. In addition to the *KALRN*-PT mutation identified in a Japanese population, private damaging *KALRN* mutations were also associated with schizophrenia in a cohort of Xhosa individuals, a southern African population with high genetic diversity (14,16). Mutations in *TRIO*, a paralog of *KALRN*, and 16p11.2 copy number variants are both among the most penetrant genetic risk factors for schizophrenia, and are implicated in cytoskeletal pathways such as RhoA regulation (15,58,59). Specifically, a bipolar disorder-linked mutation in the second GEF domain of *TRIO,* homologous to the second GEF domain in *KALRN*, was found to confer RhoA gain of function much like *KALRN*-PT (19). Mutations in several other cytoskeletal regulatory genes (GTPases, GEFs, and MAIs) were also significantly associated with schizophrenia risk in the recent large-scale analysis from the SCHEMA consortium (15,60).

The impacts we identified downstream of *Kalrn*-PT may therefore represent phenotypes driven by multiple possible dysregulators of RhoA or cell structure broadly. Given that *in vitro* evidence indicates an overactivation of RhoA by *Kalrn*-PT (with no effect on Rac1), RhoA gain of function presents a plausible molecular mechanism linking morphological and functional alterations. Nonetheless, future work should aim to verify the impacts of RhoA dysregulation on mismatch responses through i) RhoA inhibition in *Kalrn*-PT mice and/or ii) similar testing in other mouse models of RhoA dysregulation (e.g., (61,62)) to determine whether the co-occurrence of dendrite structure and mismatch response deficits can be directly attributed to RhoA overactivation as a phenotypic convergence point.

One advantage of the *Kalrn* mutation employed in this study lies in the spatially restricted phenotypes observed. Along with our past work in the auditory domain, we report that altered structure and function in *Kalrn*-PT mice is primarily observed in supragranular neurons, which may sit upstream of other impacted brain regions in schizophrenia. Supragranular neurons, which have been dubbed “comparators of top-down and bottom-up input,” underlie important perceptual functions, including context processing (i.e., deviance detection) and sensorimotor integration (34). Interestingly, cortical *Kalrn* expression skews towards supragranular cortical layers and glutamatergic neurons, providing an early mechanistic explanation for the spatially restricted phenotypes observed in *Kalrn*-PT mice (63–65).

Clinical and preclinical studies often focus on superficial layers without taking measures from other layers, making it difficult to identify which phenotypes are restricted to these layers. Our data show a single genetic mutation capable of recapitulating multiple schizophrenia-relevant phenotypes *specifically* in upper cortical layers. These layer-specific phenotypes offer an important site of convergence, within or across clinical subtypes. Further clarifying the laminar specificity of phenotypes or even biotypes may direct the development of improved therapeutics.

Previously, we established a developmental trajectory of dendritic regression across adolescence in *Kalrn*-PT mice (12). We found that deviance detection responses were heavily reduced in *Kalrn*-PT mice, suggesting a potential interaction between typical functional development in adolescence (i.e., the development of deviance detection) and dysregulated cell morphology in development. In line with this and in contrast to WT development, we found no significant differences in deviance detection between pre-adolescent and adult *Kalrn*-PT mice (Supplemental Fig. 5). This “reductionist” approach utilizing a single point mutation allows for a firmer, though of course not definitive, understanding of how typical adolescent development of cortical function relies on appropriate *Kalrn* function and, potentially, appropriate dendritic arborization.

More broadly, our results suggest that *Kalrn*-PT mice exhibit deficits in cortical integration reliant on higher order feedback. Given the reliance of deviance detection on prefrontal input, we investigated the integrity of front-visual circuitry in *Kalrn-*PT mice (31). We report a reduction in peak fronto-visual synchrony in these mice, implying blunted interregional signal integration. Our past work indicates that ACa activity drives this theta-band peak in synchrony, suggesting that V1 integration of top-down inputs may be impaired in *Kalrn*-PT mice (35). On the other hand, *Kalrn*-PT mice do not exhibit broader sensory cortical processing deficits as typical firing rates and orientation selectivity were observed in these mice (Fig. 3Q, 3R).

These functional deficits may also be limited to cortical computations. We found no genotype differences in stimulus-specific adaptation (Supplemental Fig. 3), a repetition suppression signal inherited from the thalamus. Moreover, the lack of change in orientation selectivity (Fig. 3R), which is partially present in the thalamus in mice, further supports this notion (69). These findings concur with our past data showing aberrant gap pre-pulse inhibition responses but typical noise pre-pulse inhibition responses in *Kalrn*-PT mice, indicating impaired cortical auditory processing and intact subcortical auditory processing, respectively (12).

While gross neuroanatomical changes and mismatch negativity have been correlated in individuals with schizophrenia (28–30), to our knowledge, we are the first to show a direct link between a cytoskeletal regulatory gene, dendritic morphology, and deviance detection. Other studies have identified altered auditory oddball responses in mice missing one or both alleles for a gene relevant to neuronal structure broadly (*Akap11* (70), *Src* (71), and *Nrg1* (72); however, none of these distinguished between (subcortically emergent) stimulus-specific adaptation and (cortically emergent) deviance detection.

This study specifically connects co-occurring structural and function phenotypes via the *Kalrn*-PT mutation. However, more direct mediators relating dendritic reductions and deviance detection are likely and worth further investigation. Beyond the tenable role for RhoA dysregulation noted above, it is feasible that reported alterations in supragranular pyramidal arbors can act as an immediate substrate for the loss of deviance detection and fronto-visual synchrony, as the extent of dendritic arbors can directly influence dendritic integration (38). Regardless of the direct mediators, our results suggest that typical *Kalrn* functioning is necessary to compute typical mismatch responses in supragranular pyramidal neurons and advances efforts to disentangle the specific site of convergence (i.e., the farthest downstream effect that will suffice to generate these phenotypes) for these two highly replicated phenotypes.

Altogether, our data map a disruption of *Kalrn* onto a circumscribed set of coinciding disease features, supporting a relationship between dendritic arborization and sensory mismatch responses, particularly in superficial cortical layers. While additional work is needed to clarify the precise intermediate mechanisms linking RhoA, dendritic deficits, and reduced mismatch responses, our findings present foundational evidence linking these phenotypes and underscore the value of the *Kalrn*-PT model – and other rare genetic variants – for investigating how certain phenotypes functionally connect in disease populations by cutting through the “noise” of clinical heterogeneity.

## Acknowledgements

This work was funded by F31EY036279 (A.M.R.G.), R01MH132586 (M.J.G), R01EY033950 and R01MH128176 (J.P.H). Confocal imaging was supported by S10OD032336. The authors gratefully acknowledge the GSU Imaging Core for their assistance and thank Dr. Georgia Bastos and Antanovia Ferrell for their insight and support. The content is solely the responsibility of the authors and does not necessarily represent the official views of the National Institutes of Health.

