## Supplemental Methods and Figures for "A Key Regulator of Dendritic Morphology in Supragranular Neocortex Impacts Mismatch Negativity"

*Animals.* Adult (12 weeks or older) female and male mice were used for these experiments. Mice were weaned between postnatal day (p)21 and p25 and separated by sex (maximum five mice per cage). Animals were kept under a 12-hour light/dark cycle with food and water available *ad libitum*.

Electrophysiological data for this manuscript consisted of two main experimental cohorts generated on independent experimental set-ups: Cohort 1, bipolar electrodes implanted in V1 and ACa, 12-18 weeks, n=8 WT (3 female) and n=8 *Kalrn*-PT (4 female); Cohort 2, multielectrode implanted in V1, 18-20 weeks, n=4 WT (3 female) and n=5 *Kalrn*-PT (3 female). Differences between these cohorts are described below using this notation, where applicable.

*Golgi Histology and Imaging.* Mice (*Kalrn*-PT and littermate/cagemate WT) were anesthetized and rapidly decapitated, and brains were extracted. Golgi staining was conducted using the FD Rapid GolgiStain Kit according to the associated protocol (FDNeurotechnologies). Estrus phase (follicular vs luteal) was balanced across genotype. Mice were ~12 weeks or ~18 weeks to reflect the age range recorded in Cohort 1; no qualitative differences were observed between ~12-week and ~18-week animals.

Golgi-stained tissue from primary visual cortex (V1) of WT (n=13 mice, 8 female) and *Kalrn*-PT mice (n=13 mice, 8 female) was imaged under a 30x silicon oil objective (NA 1.05) on an Olympus IX73 microscope. Z-stacks were acquired every 1.5um steps through the full thickness of the neuron. Tiling using cellSens was conducted across the entire section under 4x to determine the boundaries of V1 and gauge soma depth. A secondary scorer validated pyramidal morphology on a case-by-case basis. The basilar arbors of Golgi-stained pyramidal neurons were then manually reconstructed using the NeuroLucida software, and Sholl analyses for intersections and dendritic length were carried out for Sholl radii every 20um using NeuroExplorer.

Additional tissue (n=4 per genotype, from animals used in immunohistochemistry experiments) underwent Nissl staining (dehydration, cresyl violet immersion, differentiation, and clearance by xylenes). Sections were manually scored for laminar thickness as a percentage of total cortical thickness, and these percentages were used to assign laminar positions of Golgi-stained neurons (see Fig. 1b).

*Immunohistochemistry (IHC) and Confocal Imaging.* WT (n=4, 2 female) and Kalrn-PT mice (n=4, 2 female) were transcardially perfused with 4% paraformaldehyde (PFA) at postnatal day 84 or 85 (12 weeks), and brains were extracted, post-fixed in PFA overnight, then immersed in 30% sucrose for cryoprotection. Brains were then sectioned coronally at 25μm, and sections containing V1 or ACa were placed free-floating in 1X phosphate-buffered saline (PBS). Sections were washed in PBS (3x) then incubated for twenty minutes in 0.09% hydrogen peroxide to block endogenous peroxidase activity. Sections were washed again in PBS (3x) and moved into blocking buffer (5% normal donkey serum, 0.2% bovine serum albumin in 0.1% PBS-TritonX) for one hour to reduce background signal. Primary antibody incubation (PSD-95 hosted in rabbit (1:1000; Invitrogen 51-6900)) was done at 4°C for 72hrs, with 30 minutes of acclimation at room temperature at the start and end of the incubation. After primary incubation, the sections were washed in 0.1% PBS-TritonX (3x). Sections were then incubated in secondary antibodies (donkey anti rabbit -568 (1:500; Jackson Immuno 711-575-152)) for 2-3 hours. Sections were washed again in 0.1% PBS-TritonX and then mounted on Superfrost Plus slides and sealed with DAPI (Vectashield, Vector Laboratories).

Images were acquired on a Zeiss LSM 980 with Airyscan 2 (laser lines: 568) under a Plan-Apochromat 63x/1.40 objective with 1.8x digital zoom (74.8x74.8um ROI) in 5um z-stacks. 6 ROIs were acquired in V1 and were centered roughly around supragranular layers. Density measures were normalized by IHC round (using WT averages within each round).

*Analysis: Histology.* To assess genotype differences in dendritic morphology, the area under the curve across the 30-70um Sholl radii (proximal AUC) was taken for dendritic length and number of Sholl intersections, as proximal dendrites showed the greatest genotype difference qualitatively. A linear mixed effects model was fit to the morphological variable (proximal AUC for length or intersections) with cortical layer and genotype as fixed effects and mouse as a random effect (Y ~ Layer * Genotype + (1|Mouse)). We computed estimated marginal means (i.e., least-squares means) for analysis of genotype within cortical layer, using one-sided testing given our strong *a priori* hypotheses based on previous observations in A1 of *Kalrn*-PT mice. We also conducted a likelihood ratio test comparing the model with and without the interaction term to assess a genotype-by-layer interaction. These tests revealed no significant interaction terms, although the interaction term for dendritic length approached significance (length: X2 (2, N = 172) = 4.888, p=0.087; intersections: X2 (2, N = 172) = 3.2863; p=0.1934). P values and degrees of freedom were computed via the lmerTest (1) and emmeans (2) packages in R Studio using Satterthwaite's approximation.

To assess differences in PSD-95 density, confocal images were imported into FIJI and z-projected across 8 optical sections (selected for quality). Images were then processed using the SynBot plugin and manually thresholded to quantify puncta (3). A generalized linear model was fitted for the data with genotype as the predictor using cluster bootstrap resampling at the mouse level (ClusterBootstrap package in R; (4)). Bootstrapped samples (1,000 iterations) were generated at the mouse level to preserve the hierarchical structure, accounting for within-mouse correlations. Statistical inference was conducted via permutation testing to compare genotypes (10,000 permutations); the p value reported represents the proportion of permuted test statistics as or more extreme than the observed statistic. A one-tailed test was used given our directional *a priori* hypothesis reflecting previous dendritic spine density data in *Kalrn*-PT A1 (5).

For all histological data processing and analysis, the experimenter was blinded to genotype and the same experimenter conducted data processing and analysis to ensure consistency.

*Electrophysiology: Surgery and Training*. Animals were anesthetized with isoflurane (3% at induction, 0.8-2% for maintenance). For both cohorts, a titanium headplate was secured to the skull to allow for head fixation during recordings.

*Cohort 1 (bipolar electrodes in V1):* Prior to the surgery, two pairs of bipolar titanium electrodes (Plastics One, Roanoke, VA, USA) were twisted together. Mice were implanted with one twisted bipolar electrode inserted below the dura in stereotaxically defined left V1 (coordinates from bregma: V1, X=2 mm, Y= -2.92 mm) and a second twisted bipolar electrode was implanted in left anterior cingulate area (ACa; X = -0.35 mm, Y = +1.98 mm, Z = 0.9mm). Bipolar electrodes were grounded on the contralateral skull.

*Cohort 2 (multielectrode in V1):* In the first surgery, a single titanium reference screw overlying the cerebellum was also implanted at the time of headplate attachment. A second surgery was performed one day prior to the experiment creating a small craniotomy (1-2 mm) above left V1 for acute electrode insertion. Prior to insertion during surgeries, probes and electrodes were submerged in DiI dye for post-hoc anatomical validation.

Following surgery, each animal received three days of head fixation training on a manual treadmill, increasing in duration from 5-10 minutes to 45-60 minutes, to improve tolerability and comfort during head fixation. During training, mice were exposed to sequences of visual moving grating stimuli (described below) for acclimation.

After recordings, brains were post-fixed over-night in 4% paraformaldehyde, then cryosectioned coronally and visually inspected for electrode placement confirmation, referenced against the Allen Brain Atlas. For Cohort 2, 3D reconstructions of multielectrode tracks were done using SHARP-Track for registration to the Allen CCF (6).

*Visual Stimuli.* As previously described (7,8), full-field square-wave gratings (100% contrast; .08 cycles per degree) of 22.5, 45, 67.5, 90, 112.5, 135, 157.5, and 180-deg orientations were created with MATLAB Psychophysics Toolbox (9,10) and displayed on an LCD monitor (19-inch diameter, 60 Hz refresh rate for Cohort 1, 27-inch diameter, 170 Hz refresh rate for Cohort 2) positioned ≈20-30cm from the right eye while the mouse was head-fixed and freely moving on a treadmill.

Animals were exposed to two types of visual sequences. i) The “many-standards” control sequence is made up of all 8 orientations presented randomly with approximately equal likelihood (≈12.5% probability) (250 trials in Cohort 1, 400 trials in Cohort 2). ii) The oddball sequence run is composed of two stimuli separated by 90-deg (e.g., 0-deg and 90-deg), one shown in a redundant context (≈87.5% probability) and one in a deviant context (≈12.5%) presentation (125 trials for Cohort 1, 200 trials for Cohort 2); contexts were “flip-flopped” within the same run, halfway through, such that the previously redundant stimulus served as the deviant and vice versa (125 more trials for Cohort 1, 200 more trials for Cohort 2).

For Cohort 1, experiments consisted of alternating control and oddball sequence runs (two of each), whereas in Cohort 2, two oddball runs followed one control run. In the second oddball run, a different stimulus orientation pair (e.g., 45-deg and 135-deg) was shown. This procedure ensures that four separate orientated stimuli participated in all three sensory contexts: equiprobable (control), high likelihood (redundant), and rare (deviant). Visual trials across all paradigms were 1.025 second in duration: 500ms of drifting gratings interleaved with 500-550ms of a middle gray blank screen.

*Electrophysiology: collection and processing.* Intracortical electrical signals were recorded from either a) bipolar electrodes inserted into V1 and ACa (Cohort 1) or b) custom designed 16-channel NeuroNexus probe (750µm length, 50µm inter-contact distance; A1×16–3mm50–177; Ann Arbor, MI) inserted perpendicularly into left V1 at 1µm/s until the dorsal-most electrode was 25-50µm below the dura (deduced from real-time signals; Cohort 2).

*Cohort 1:* Insulated cables were connected to the implanted electrodes (ACa and V1) and plugged into a differential amplifier (Warner instruments, DP-304A, high-pass: 0 Hz, low-pass: 500 Hz, gain: 1K, Holliston, MA, USA). Amplified signals were passed through a 60 Hz noise cancellation machine (Digitimer, D400, Mains Noise Eliminator, Letchworth Garden City, UK), which, instead of filtering, creates an adaptive subtraction of repeating signals which avoids phase delays or other forms of waveform distortion.

*Cohort 2:* Multielectrode signals were amplified and digitized at 20kHz using an Intan 512ch recording controller (0.1 Hz lower bandwidth; 7500 Hz upper bandwidth) after a minimum of 20 minutes elapsed following insertion.

*Analysis: single contact bipolar electrode recordings.* All analyses were performed using MATLAB (MathWorks 2020) using code written in house and publicly available (11). For quantifying local field potentials (LFP) in V1 (Cohort 1; n=8 *Kalrn*-PT, n=8 WT), data were analyzed in the time-frequency domain using EEGLAB (12) to apply modified Morelet-wavelets to LFP data from individual trials, comprising 100 equally-spaced wavelets (4-80Hz, 1Hz resolution, linearly increasing from 1 cycles to 20 cycles) applied every 10ms from -500ms pre-stimulus onset to 500ms post stimulus offset. Stimulus-induced power was quantified as the across-trial average of the squared magnitudes of the absolute value of the complex output of the wavelet analysis. For all analyses, we used the 100ms pre-stimuli as a baseline and subtracted values on a trial-wise, electrode-wise basis. These data were combined with the multicontact probe for LFP power comparisons (see below).

For ACa-to-V1 coherence in the two-bipolar electrode group (Cohort 1; n=7 WT, n=8 *Kalrn*-PT), we computed the phase difference between ACa and V1 vectors for each frequency for each 10ms time bin (-250ms to 750ms relative to stimulus onset) and trial. This phase difference was assigned a unit amplitude to generate a complex phasor, and then phasors were averaged across timepoints and trials within each frequency. The absolute value of this result was bound between 0 and 1, with 1 reflecting perfectly consistent phase lags between V1 across time (perfect phase locking) and 0 reflecting no locking at all.

*Analysis: multielectrode recordings.* For analysis of stimulus-induced power in the multielectrode recordings, we repeated the above wavelet analysis on bipolar electrode contacts from Cohort 1. Effects were nearly identical across layers (as LFP has relatively low spatial resolution compared to single units), so we collapsed across contacts within each mouse. Trial numbers were equated across conditions (control and deviant) within mice. We combined single trial power estimates across both methods (single and multielectrode contexts) and focused on theta to low-gamma power (4-60Hz), averaged from 40 to 350ms post stimulus onset and across orientations, producing one value for each mouse and context for statistical analyses. We carried out a linear mixed effects analysis with context as a within-subjects variable and genotype and sex between-subjects variables. Mouse and cohort (single vs multielectrode recordings) were treated as a random effects variable.

For analysis of neuronal spiking, we analyzed single and multiunit activity in multielectrode recordings. Preprocessing for spike sorting was carried out by first concatenating time-series signals from many-standards control and oddball paradigm recordings, converting to 16-bit signed integer, high-pass filtering at 300 Hz, and taking the common average reference (13) to remove spatially diffuse artifacts. Signals were then Zero-phase component analysis (ZCA) whitened to spatially filter and de-correlate the channels. Single and multiunit clusters were identified as threshold crossings exceeding 5 standard deviations over 12-30s batches, and their spike times and waveforms extracted using Kilosort4 (14). Resulting Kilosort4 outputs were then manually curated in Phy (15) where isolation quality was assessed through inspection of waveform stereotypy, refractory period violations (i.e. false positive rate) and the separability of clusters in feature space. Waveforms were then extracted from high-pass filtered data in 3ms windows (1.5ms on either side of extracted spike times) centered on the trough. Eighty-three units satisfied the most stringent criteria and represented stable, well-isolated single units. An additional 101 units showed otherwise strong stability but had wider variance in their waveform properties and, thus, may have reflected either isolated neurons or multineuron clusters. Such clusters were combined with single units for the analysis of visual context, as deviance detection has been shown in both single neuronal (16) and multiunit (7) responses. The same pattern of effects (deviance detection, orientation selectivity, firing rates) was present when analyses were limited only to well-isolated single units.

Units were segregated into broad spiking and narrow spiking units/clusters based on trough-to-peak duration (the interval between the global minimum and following local maximum) and a repolarization time (interval between late positive peak and inflection point of falling flank) as in previous works ((17); See Supplemental Fig. 2).

For each unit we averaged spikes across trials in which mice saw a given orientation as a contextual deviant (during the oddball) or contextually "neutral" control (n=15-25, equated across conditions within recordings). After averaging, we smoothed firing rates (FR) with a 25ms moving mean window and normalized using the standard deviation of activity across all conditions (Norm FR). For neurons exhibiting significant orientation selectivity (during the many-standards control, 1-circ variance > 0.1; 38.1% total), we only used one orientation in the main analysis (i.e. the maximum). Otherwise, we averaged across both orientations. Orientation selectivity index (OSI) was calculated using the many-standards control runs as 1-circular variance, a more conservative, albeit reliable, approach for estimating feature selectivity in cortical recordings (18).

For statistical analyses of deviance detection, we focused on the early stimulus evoked peak (occurring in the first 150ms post stimulus onset) and the average sustained activity (150ms-350ms) separately. For analysis of stimulus specific adaptation, we used the third redundant after each deviant in order to equate number of trials across conditions, and we focused on the early mean evoked activity in the first 100ms post-stimulus based on our past work with extracellular electrophysiology (7,19).

**Supplemental Figures**

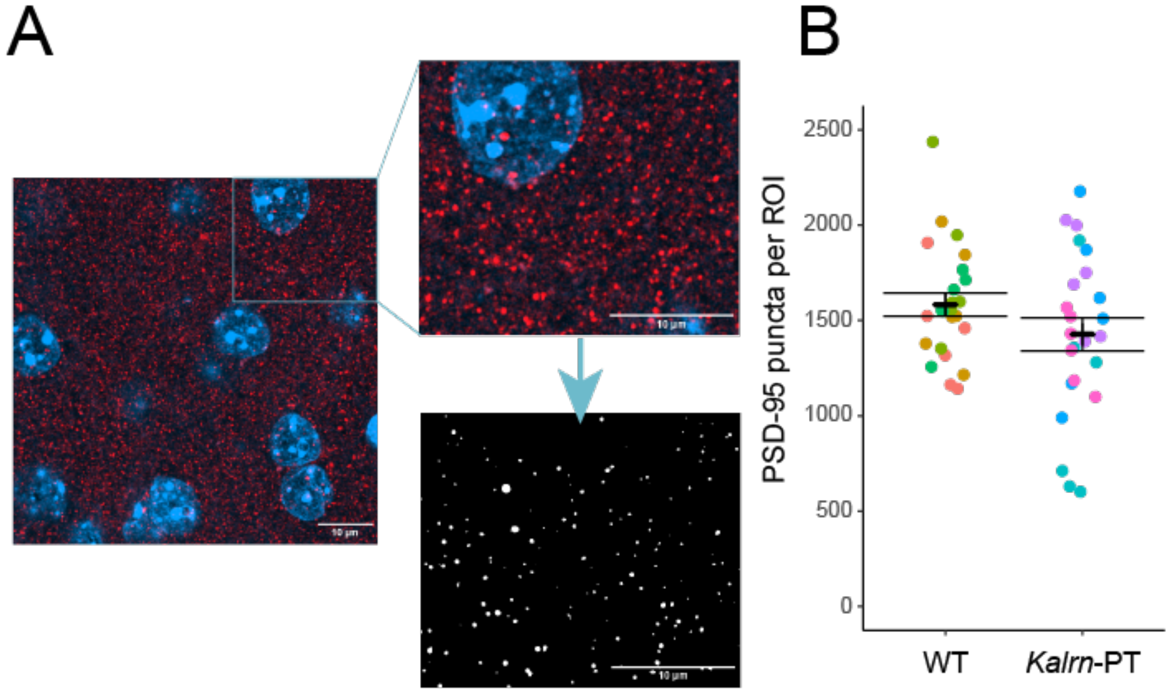


**Supplemental Figure 1. *Kalrn*-PT mice exhibit a trend towards decreased V1 PSD-95 density.**

**(A)** Immunostained PSD-95 puncta (red) and DAPI (blue) in a 67x67um ROI from V1, max projected across 8 Z planes sampled every 0.15um (left). The inset (right) illustrates the thresholding approach using the Synbot FIJI plug-in to identify puncta. Enlarged to show detail. **(B)** A non-significant trend towards reduced PSD-95 density (9.9% decrease) in superficial V1 (t=1.459, p=0.076) follows the pattern previously reported in *Kalrn*-PT A1. The t statistic was determined from permutation testing (10000 permutations) on clustered bootstrap samples (1000 iterations) generated at the mouse-level to account for within-mouse correlations. One-sided tests were conducted given our *a priori* expectations based on previous data in A1 (5). n=4 mice per genotype, 6 ROIs per animal. Matching colored dots correspond to ROIs from the same animal. Error bars indicate standard error of the mean.

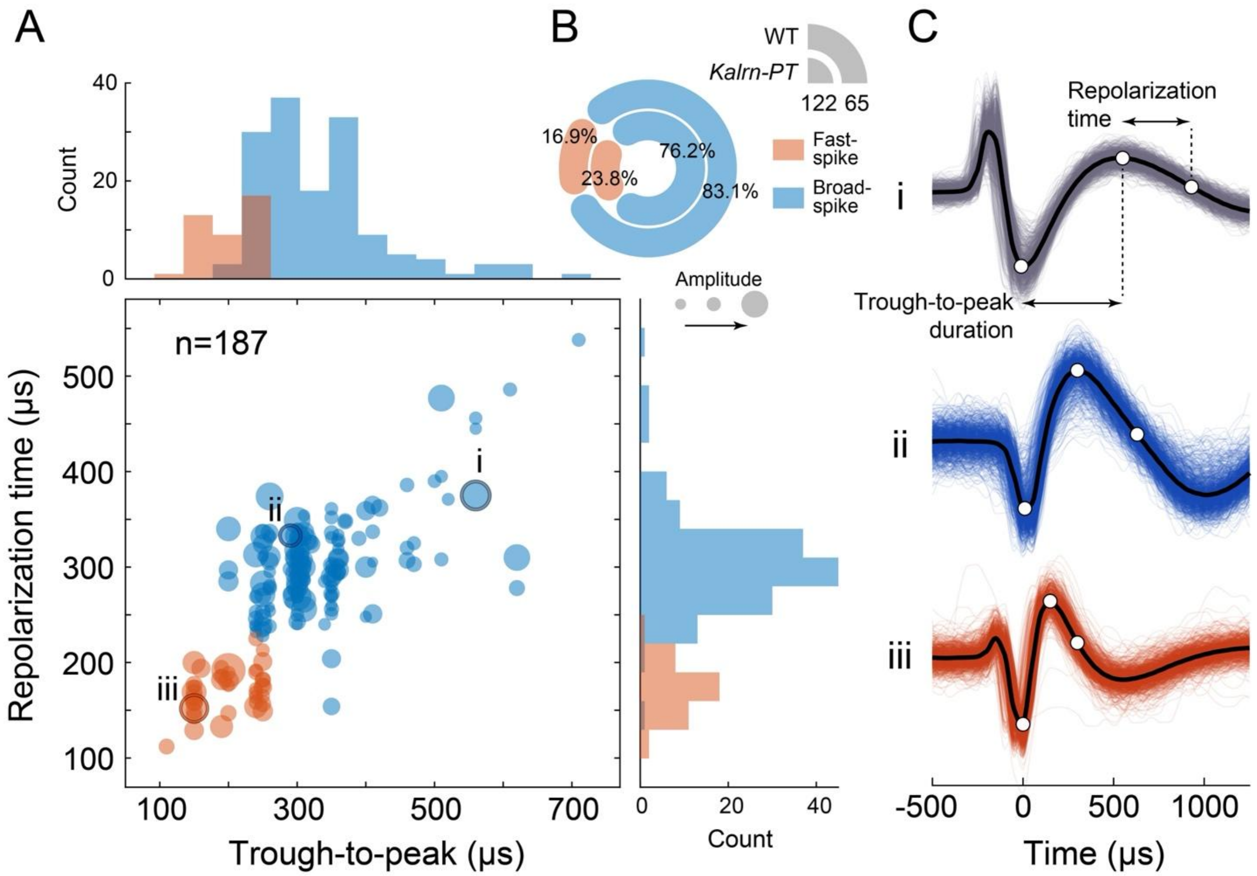


**Supplemental Figure 2. Unit isolation and segregation by group.**

**(A)** Scatter plot of discriminative waveform characteristics and marginal distributions for all single and multiunit clusters recorded in this study (definitions illustrated in (C)), with dot size scaled by the unit's amplitude. **(B)** Relative proportions of fast- and broad-spiking waveforms by Kalrn-PT and wild-type (WT) genotypes, with total counts inset below the radial legend. **(C)** Representative waveforms (n=1000) for well-isolated single unit clusters (i-iii) in (A) ranging from broad- to fast-spiking, respectively. Average waveforms are in black. White circles correspond to the trough, peak, and inflection point indicating half the decay amplitude.

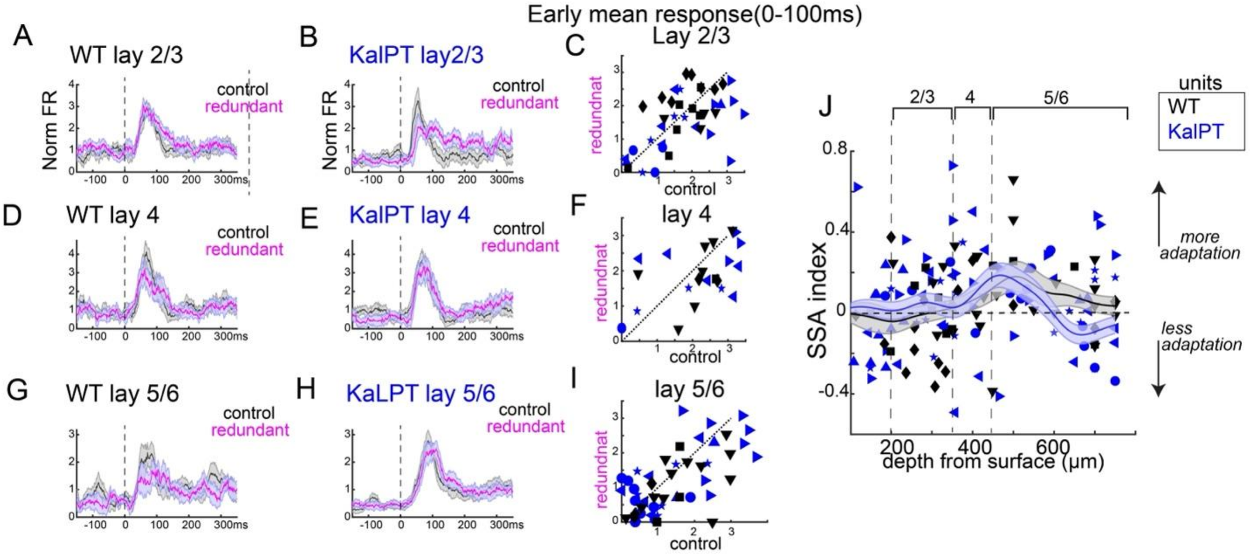


**Supplemental Figure 3. Stimulus specific adaptation to redundant stimulus is not altered in *Kalrn*-PT mice.**

**(A)** Responses to the same orientation when it was contextually neutral vs contextually redundant, averaged over broad spiking units from wild-type mice (shaded area indicates standard error of the mean). **(B)** Same as A for *Kalrn*-PT animals. **(C)** Mean responses from 0 to 100ms after the onset of stimuli. Each dot is one neuron. **(D-F)** Same as A-C for layer 4 neurons. **(G-I)** Same as A-C for layer 5/6 neurons. **(J)** Stimulus specific adaptation index (control minus redundant, divided by max response over all contexts) plotted as a function of depth from surface. Smoothed gaussian fit line (200µm window) for each group. Shapes correspond to units from the same animal.

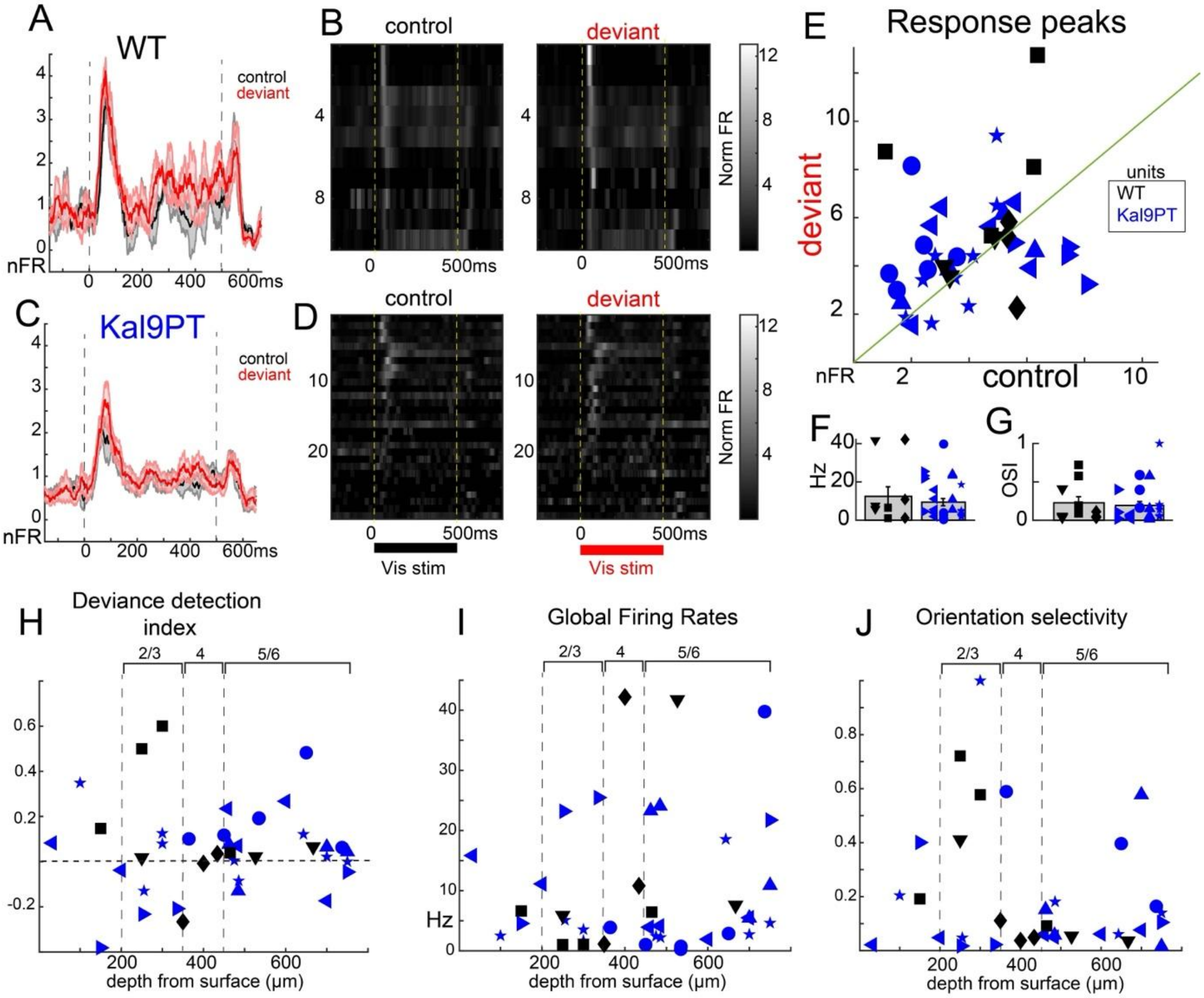


**Supplemental Figure 4. Narrow spiking units are not altered in *Kalrn*-PT mice.**

**(A)** Trial averaged responses of narrow spiking units to deviant vs control stimuli across units and **(B)** presented for individual units in WT mice. **(C-D)** Same as A-B for *Kalrn*-PT mice. **(E)** Scatter plot of peak responses for each unit to the control vs deviant, as in main figure 3E, J, O, all layers included. Shapes correspond to units from the same animal. No group differences were found globally (t(74)=-0.88, p=0.35) or for any individual layer. **(F)** Global (all recordings) firing rates did not differ (t(37)=0.70, p=0.48), and **(G)** Orientation selectivity (1 minus circular variance) did not differ (t(31)=0.37, p=0.71). **(H-J)** Values from E-G, plotted as a function of layer.

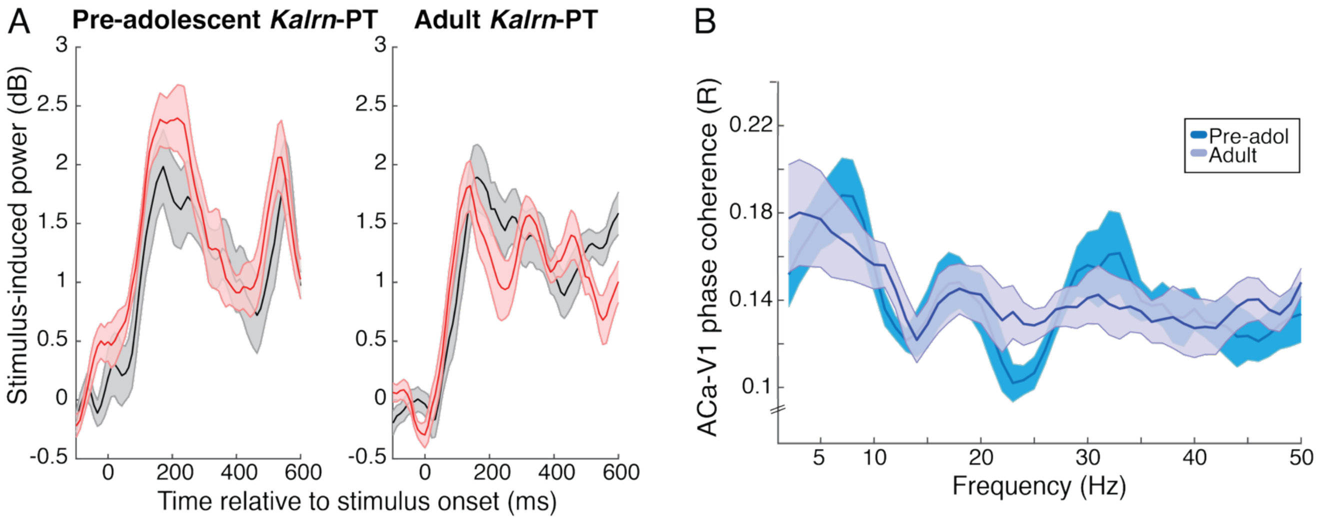


**Supplemental Figure 5. Deviance detection fails to develop in *Kalrn*-PT mice.**

**(A)** Stimulus-induced responses by context (± standard error of the mean) for pre-adolescent (n=7; p28-p31) and adult (n=8; Cohort 1) *Kalrn*-PT mice during V1 bipolar electrode recordings revealed no differences across development ($F(1,13)=1.16$, $p=0.30$). **(B)** ACa-V1 phase coherence (± standard error of the mean) during visual stimulation, averaged across all stimulus conditions also did not differ between pre-adolescent (blue) and adult (purple) *Kalrn*-PT animals ($F(1,13)=0.57$, $p=0.46$).

numbers into movies. *Spat Vis* 10: 437–442.
